# RsQTL: correlation of expressed SNVs with splicing using RNA-sequencing data

**DOI:** 10.1101/840504

**Authors:** Justin Sein, Liam F. Spurr, Pavlos Bousounis, N M Prashant, Hongyu Liu, Nawaf Alomran, Jimmy Bernot, Helen Ibeawuchi, Dacian Reece-Stremtan, Anelia Horvath

**Affiliations:** McCormick Genomics and Proteomics Center, School of Medicine and Health Sciences, The George Washington University, 20037 Washington, DC, USA; Department of Medical Oncology, Dana-Farber Cancer Institute, Boston, MA 02215; Cancer Program, The Broad Institute of MIT and Harvard, Cambridge, MA 02142; Computer Applications Support Services, School of Medicine and Health Sciences, The George Washington University, 20037 Washington, DC, USA; Department of Biochemistry and Molecular Medicine, Department of Biostatistics and Bioinformatics School of Medicine and Health Sciences, George Washington University, 20037 Washington, DC; Department of Pharmacology and Physiology, School of Medicine and Health Sciences, The George Washington University, 20037 Washington, DC, USA

## Abstract

RsQTL is a tool for identification of splicing quantitative trait loci (sQTLs) from RNA-sequencing (RNA-seq) data by correlating the variant allele fraction at expressed SNV loci in the transcriptome (VAF_RNA_) with the proportion of molecules spanning local exon-exon junctions at loci with differential intron excision (percent spliced in, PSI). We exemplify the method on sets of RNA-seq data from human tissues obtained though the Genotype-Tissue Expression Project (GTEx). RsQTL does not require matched DNA and can identify a subset of expressed sQTL loci. Due to the dynamic nature of VAF_RNA_, RsQTL is applicable for the assessment of conditional and dynamic variation-splicing relationships.

**Availability and implementation:** https://github.com/HorvathLab/RsQTL.

**Supplementary Information:** RsQTL_Supplementary_Data.zip

## 1. Introduction

Splice QTLs (sQTLs) are involved in phenotype formation and complex disease risk, to a level comparable or higher to that of eQTLs (Li, Y. I. *et al*, 2016, Brandt M., and Lappalainen T, 2017). eQTLs and sQTLs are both traditionally assessed from matched DNA and RNA datasets, where DNA is used for genotype estimation and RNA for expression (eQTLs), or splicing (sQTLs) estimation. We have recently developed a method to assess eQTLs from RNA-seq data alone-ReQTL (Spurr, L. *et al*, 2019) - which replaces the genotypes with the variant allele fraction, VAFrna, and identifies a subset of eQTLs. We present a related method, RsQTL (**R**NA-**sQTL**), which identifies splicing QTLs via correlation of VAFrna with the proportion of excised introns (percent spliced in, PSI) at loci with differential intron excision (Li, Y. I. *et al*, 2018).

We demonstrate RsQTL using Matrix eQTL (Shabalin, 2012) on RNA-seq data obtained from the Genotype-Tissue Expression (GTEx) project (www.gtexportal.org, phs000424.v7), from three different tissue types: Nerve-Tibial (NT), Skin-Sim-Exposed (SkE), and Skin-Not-Sim-Exposed (SkN). The proposed pipeline (Figure 1a, S_Table 1, and S_Methods) employs publicly available packages for processing of RNA-seq data and a toolkit for RsQTL-specific data transformation (https://github.com/HorvathLab/RsQTL). RsQTL analyses are optimized for SNV (Single Nucleotide Variants)-aware alignments, produced via a two-pass alignment strategy (STAR, v.2.7.2, Dobin, A., *et al.*, 2013). Briefly, SNVs are called from the non-SNV-aware alignments, (GATK v.4.0.8.0, Van der Auwera, G.A. *et al.* 2013) and combined into a list of unique positions, which are then inputted into WASP (Van de Geijn, B, *et al.*, 2015) to correct for allele mapping bias during the second alignment. These SNV-aware alignments are then used to estimate: (1) VAF_RNA_ using ReadCoimts (Movassagh, M *et al.*, 2017), and (2) PSI using LeafCutter (Li, Y. I. *et al*, 2018). VAF_RNA_ was estimated from loci covered by a minimum of 10 RNA-seq reads, and PSI was estimatedfrom intron clusters covered by a minimum of 30 RNA-seq reads. VAF_RNA_ and PSI are then combined into matrices, and filtered to remove SNV- and intron-loci not informative or homozygous across more than 80% of the studied individuals. Principal components (PC) are computed to account for hidden confounders and used as covariates (together with known covariates). The VAF_RNA_, PSI and covariate matrices are then used as inputs for Matrix_eQTL; RsQTLs are assessed using a linear regression model and a false discovery rate (FDR) of 0.05. A parallel sQTL analysis is performed to assess overlapping and exclusive outcomes.

**Figure 1.**
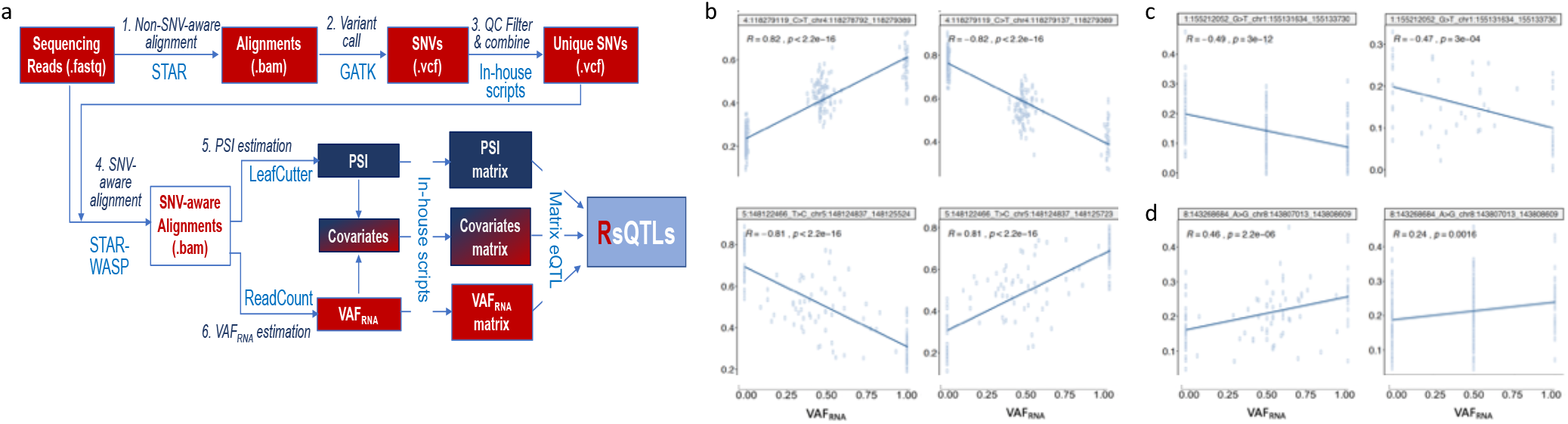
**a.** RsQTL analyses (differences from eQTL analysis are outlined in red). SNV-aware alignments are generated using STAR (two-pass strategy), where the SNVs called on the 1^st^-pass alignments are used (1) by WASP to remove ambiguously aligned reads, and, (2) by ReadCounts to estimate VAFRNA from the 2^nd^-pass (SNV-aware) alignments. Note that the latest versions of STAR have the WASP-option implemented, which streamlines the process significantly. The same SNV-aware alignments were used to estimate PSI. VAFRNA and PSI matrices, together with covariates, were then used as inputs to Matrix eQTL. **b.** RsQTL correlation patterns; both sQTL-like patterns (top) and patterns with VAFRNA values spread along the regression line (bottom) are seen. **c.** sQTL-exclusive correlations (left) and corresponding RsQTLs (right), NT. Genotypes were available for all the samples while VAFRNA was estimated (using the required minimum 10 reads) for only 31% of the positions, **d.** RsQTL-exclusive correlations (left) and their corresponding sQTLs (right). RsQTL correlation is stronger due to the distribution of VAFRNA along the regression line. The p-values shown on the graphs are not adjusted for multiple comparisons.

## 2. Results

### 2.1. Overall

For direct comparisons with sQTLs, we used the same input lists of SNV loci per tissue, which were generated based on accessibility for RsQTL analysis (as described above) and availability of genotypes (104054, 92776, and 94321 SNVs for the NT, SkE and SkN, respectively). Similarly, we used the same intron inputs per tissue (1710, 1387 and 1310 introns for the NT, SkE and SkN, respectively). To account for covariates, we corrected for the reported race, sex, the top three VAF_RNA_ or genotype PCs, for RsQTL and sQTL, respectively, and the top 10 PCAs for the PSI. We retained for further analyses only cis-RsQTL (SNV and intron located within 1e10^6^ nt of each other). In addition to the distance-based cis-annotation, we enable annotation based on residence in the same gene.

The numbers of significant RsQTL and sQTL-findings are shown in S_Table 2. Quantile-quantile (QQ) plots are shown in S_Figure 1, and shared and tissue-specific RsQTLs are presented in S_Figure 2. Percent explained variation by the top 10 PCs for VAF_RNA_, genotypes and PSI is shown in S_Figure 3.

Examples of significant RsQTL are shown in Figure 1b. We observed sQTL-like patterns (top), and patterns where the intermediate VAF_RNA_ values are spread along the regression line (bottom). As expected, many SNVs correlated inversely (variant vs reference nucleotide) with alternative intron excision.

### 2.2. RsQTL vs sQTL

To evaluate the proportion of sQTLs identifiable through RsQTL analysis, we analyzed overlapping and exclusive RsQTL and sQTL outputs (S_Tables 2-4). The correlations called by both methods represented 87-90% of the significant RsQTLs, and 54-57%) of the sQTLs. Accordingly, in a side-by-side setting, up to a half of the sQTLs are not called through RsQTL analyses, while approximately 10%) of the significant RsQTL correlations are not captured as sQTLs. sQTL exclusive correlations are exemplified in Figure 1c (left), together with a corresponding plot using the VAF_RNA_ from the same sample (right). On the other hand, RsQTL-exclusive correlations were observed for relatively weak sQTLs where the spread of the bi-allelic VAF_RNA_ along the regression line contributed to the detection of a stronger linear relationship (Figure 1d).

## 3. Discussion

We have previously presented a systematic analysis of VAF_RNA_ usage in QTL pipelines (Spurr, L. *et al*, 2019). Briefly, there are several important considerations for RsQTL applications. First, RsQTL analyses are confined to expressed loci and are not designed to capture SNVs in transcriptionally silent sites. Second, among RsQTL-accessible SNVs, RsQTL captures on average between 50 and 60% of the sQTL-identifiable correlations due to the lower availability of VAF_RNA_ values (as compared to genotypes). The latter is expected to be significantly improved with increased sequencing depth. Third, RsQTL might capture co-allelic (in linkage disequilibrium, LD) SNVs as opposed to the actual regulatory variant, and therefore require validation analyses before causality can be inferred. However, in comparison to ReQTL, we expect that RsQTL will capture higher proportion of causative SNVs due to the known enrichment of sQTLs within gene bodies (Li, Y. I., et al, 2016) and the related involvement of RNA-binding regulatory factors. On the other hand, due to the continuous nature of the VAFrna measure, RsQTL analyses identify about 10%o more correlations than sQTL analyses.

Several technical advantages are noted for the usage of VAF_RNA_ (Spurr, L. et al., 2019). Briefly, these include (1) the above-mentioned continuous nature of VAF_RNA_ that allows for precise quantitation of the allele counts, (2) the potential for use in identifying post-transcriptionally generated variants through processes such as RNA-editing, and (3) reduced technical noise due to estimation of both VAF_RNA_ and PSI from the same RNA-seq dataset. An additional, RsQTL-specific, advantage is that VAF_RNA_ and PSI values belong to the same interval {0,1} and therefore do not require scaling and transformation. Finally, when interpreting RsQTL results, it is important to consider the dynamics and tissue-specificity of the both VAF_RNA_ and PSI.

In conclusion, we envision applications of RsQTL for data sets where matched DNA is not available, and particularly for assessing within-gene variants which alter motifs recognizable by RNA-binding molecules. The RsQTL toolkit supports the entire pipeline from variant calls to final outputs and is accompanied by visualization modules and user-friendly instructions. All the scripts are made flexible to accommodate a large range of user-defined settings.

