## Supplementary material for "RsQTL: correlation of expressed SNVs with splicing using RNA-sequencing data": S_Methods

### Supplementary Methods

#### 1. Overall approach

We demonstrate RsQTL using Matrix eQTL (Shabalin, 2012) on RNA-seq data obtained from the Genotype-Tissue Expression (GTEx) project (www.gtexportal.org, phs000424.v7), from three different tissue types: Nerve-Tibial (NT), Skin-Sun-Exposed (SkE), and Skin-Not-Sun-Exposed (SkN). The proposed pipeline (See Figure) employs publicly available packages for processing of RNA-seq data, and a toolkit for RsQTL-specific data transformation (https://-github.com/HorvathLab/RsQTL). RsQTL analyses are optimized on SNV (Single Nucleotide Variants)-aware alignments, produced via two-pass alignment strategy (STAR, v.2.6.1c, Dobin, A., *et al*., 2013). Briefly, SNVs are called from the non-SNV-aware alignments, (GATK v.4.0.8.0, Van der Auwera, G.A. *et al.* 2013) and combined in a list of unique positions, which are then inputted in WASP (Van de Geijn,B, *et al*., 2015) to correct for allele mapping bias during the second alignment. These SNV-aware alignments are then used to estimate: (1) $VAF_{RNA}$ using ReadCounts (Movassagh, M *et al*., 2017), and (2) PSI using LeafCutter (Li, Y. I. *et al*, 2018). $VAF_{RNA}$ was estimated from loci covered by a minimum of 10 RNA-seq reads, and PSI was estimated from intron clusters covered by a minimum of 30 RNA-seq reads. $VAF_{RNA}$ and PSI are then combined into matrices, and filtered to remove SNV- and intron-loci not used or not variable across more than 80% of the studied individuals. Principal components (PC) are computed to account for hidden confounders and used as covariates (together with known covariates). The $VAF_{RNA}$, PSI and covariate matrices are then imputed in Matrix_eQTL, and RsQTLs are assessed using a linear regression model and a false discovery rate (FDR) of 0.05. A parallel sQTL analysis is performed to assess overlapping and exclusive outcomes.

#### 2. Samples

The data and analyses presented in the current publication are based on the use of study data downloaded from the dbGaP web site, under dbGaP accession phs000424.v7.p2 (Genotype-Tissue Expression (GTEx). A total of 656 raw RNA-seq datasets from three different body sites – Nerve–Tibial (NT, 197 samples), Skin-Exposed, (SkE, 243 samples), and Skin-Non-Exposed, (SkN, 216 samples) - were downloaded on 06/10/18 (S_Table 1). The samples were selected based on the availability of directly estimated genotypes (for sQTL comparisons). All the RNA-seq libraries were generated using non-strand specific, polyA-based Illumina TruSeq protocol, and sequenced to a median depth of 78 million 76-bp paired-end reads. The selection of tissue types was based on the availability of more than 150 samples with genotypes, and consideration for assessment of both distinct (NT vs Skin) and related (SkE vs SkN) tissue types.

#### 3. RNA-seq data processing

##### *3.1. Alignment using STAR-WASP pipeline*

SNV-aware alignment was performed using STAR alignment (Dobin, *et al*., 2013) followed by removal of ambiguously aligned reads using WASP (Van de Geijn *et al*., 2015). First, we aligned the RNA-seq reads to GRCh38, using STAR (v.2.6.1c) in 2-pass mode with transcript annotations from assembly GRCh38.79. We called SNVs on the alignments (see below) and combined the SNVs called across all samples from a tissue type into a list of unique SNV positions. This list was then used as an input to WASP (Van de Geijn *et al*., 2015) to test for allele mapping bias and to remove reads with ambiguous mapping due to an SNV. The generated alignments were processed for VAF and PSI estimation.

##### *3.2. Variant Call*

To call variants from RNA-seq data we used GATK (v. 4.0.8.0) and followed the provided best practices (Van der Auwera *et al*., 2013). Briefly, we first marked duplicates to clean the data, then used the module SplitNCigarReads to reformat intron-spanning reads, followed by Base Quality Score Recalibration to re-adjust the base quality values. The datasets were then subjected to variant calling using the module HaplotypeCaller. Indel calls, and mitochondrial and contig variants were filtered out. Using this pipeline, we called between 214,043 and 685,959 (average 355,201) SNVs in the individual samples from the HISAT2 alignments, and between 225,117 and 716,640 (average 371,610) from the STAR-WASP alignments. To retain high-quality SNV calls, we applied the VariantFiltration GATK module using as hard filters QUAL (Phred quality score) >100 and MQ (mapping quality) >60, and combined the filtered SNVs into a list of unique SNV positions per tissue (HISAT2/STAR-WASP: NT - 1,038,361/1,204,315, SkE – 950,858/1,076,441, SkN – 932,665/ 966,812). After annotation (SeattleSeq (v.14, DbSNP151), we retained SNVs present in the HISAT2 index, positioned outside repetitive regions, and with genotypes available from GTEx. These SNV lists were used for WASP re-alignment (see above) and for $VAF_{RNA}$ estimation and subsequent RsQTL and sQTL analyses.

#### *3.3. Variant Allele Fraction ($VAF_{RNA}$) estimation*

Within a tissue type, we estimated $n_{var}$ and $n_{ref}$ and computed $VAF_{RNA}$ ($VAF_{RNA} = n_{var} / (n_{var} + n_{ref})$) for each of the positions in the list in each of the individual samples using the module readCounts previously developed in our lab (http://github.com/HorvathLab/NGS/tree/-master/readCounts (Movassagh *et al.*, 2016). Briefly, readCounts employs the pysam Python module to assess the read counts at every SNV position of interest in each of the alignments (samples) from a studied group (i.e. tissue). ReadCounts then filters aligned reads based on alignment quality metrics including length, gaps and mapping quality, and categorizes the remaining reads as having either the reference or variant nucleotide. For RsQTL analyses, we retained only positions covered by a minimum of 10 total sequencing reads (RsQTL-fit $VAF_{RNA}$); samples with $VAF_{RNA}$ estimated from < 10 reads were assigned NA in the input matrices. Additionally, we excluded SNV positions with a monoallelic or missing (NA) signal in more than 80% of the samples from each tissue.

#### *2.2.4. PSI estimation*

Introns were clustered for each sample and their usage was quantified from the alignments using LeafCutter (version 0.2.8) (Yang I. Li, *et al*.). Introns supported by fewer than 10 reads or with a length greater than 100,000 bp were removed. Additionally, introns containing a proportion of reads smaller than 0.1 within the cluster were filtered out. 'Percent spliced in' (PSI or $\Psi$) was then estimated using read counts for each intron as a fraction of the total number of reads within the cluster. Introns containing either constitutive or unused splice junctions (PSI = 1 or 0, respectively) or insufficient data (PSI = NA) across more than 20% of samples were removed. Only introns from standard autosomal chromosomes were included in this analysis. Furthermore, within each tissue, we filtered out introns belonging to clusters covered by less than 30 sequencing reads in more than 80% of the samples. The effects of unobserved confounding variables were quantified using Principal Component Analysis, and the top 10 PCAs were used.

### 4. RsQTL analyses

https://github.com/HorvathLab/RsQTL

$VAF_{RNA}$ and PSI are combined into matrices, and filtered to remove SNV- and intron-loci not used or not variable across more than 80% of the studied individuals. Principal components (PC) are computed to account for hidden confounders and used as covariates (together with known covariates). The $VAF_{RNA}$, PSI, and covariate matrices are then imputed in Matrix_eQTL, and RsQTLs are assessed using a linear regression model and a false discovery rate (FDR) of 0.05.

### 5. sQTL analyses

A parallel sQTL analysis is performed to assess overlapping and exclusive outcomes. The genotypes for each individual were obtained from DbGaP (phs000424.v7.p2), and the PSI, covariates, and regression model were the same as those used for the RsQTL analyses. We considered significant associations after p-value correction using FDR of 5%.
