## Supplementary figures and images for "RsQTL: correlation of expressed SNVs with splicing using RNA-sequencing data"

### S_Figure_1

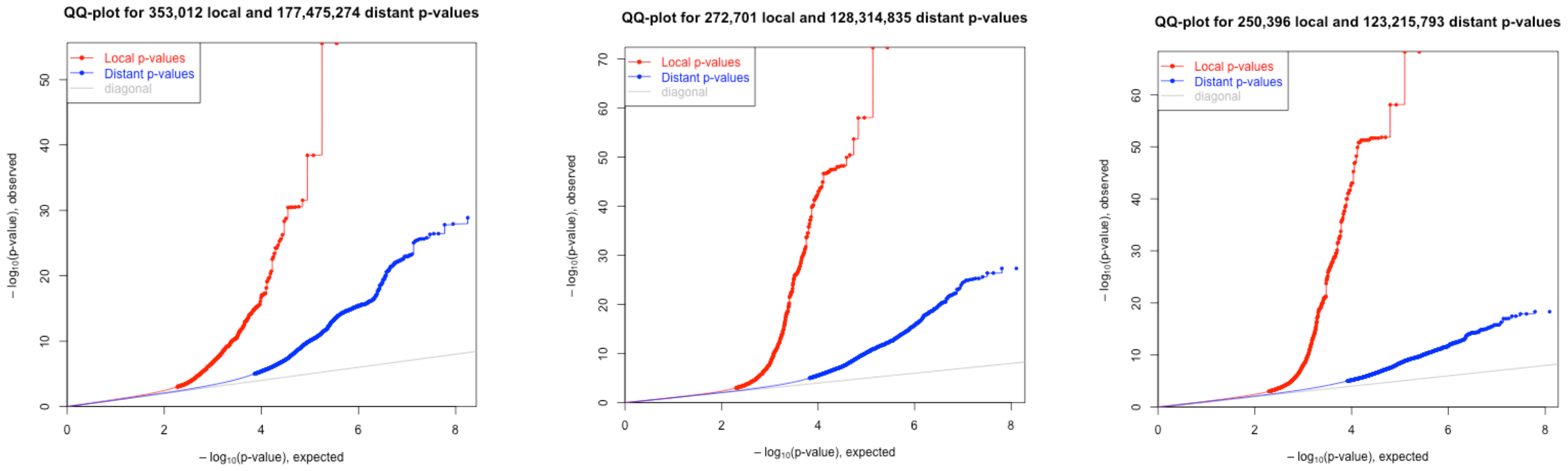


**Supplementary Figure 1.** QQ-plots of the ReQTL p-values: from left to right: NT, SkE, SkN.
