## Supplementary material for "RsQTL: correlation of expressed SNVs with splicing using RNA-sequencing data": S_Figure_2

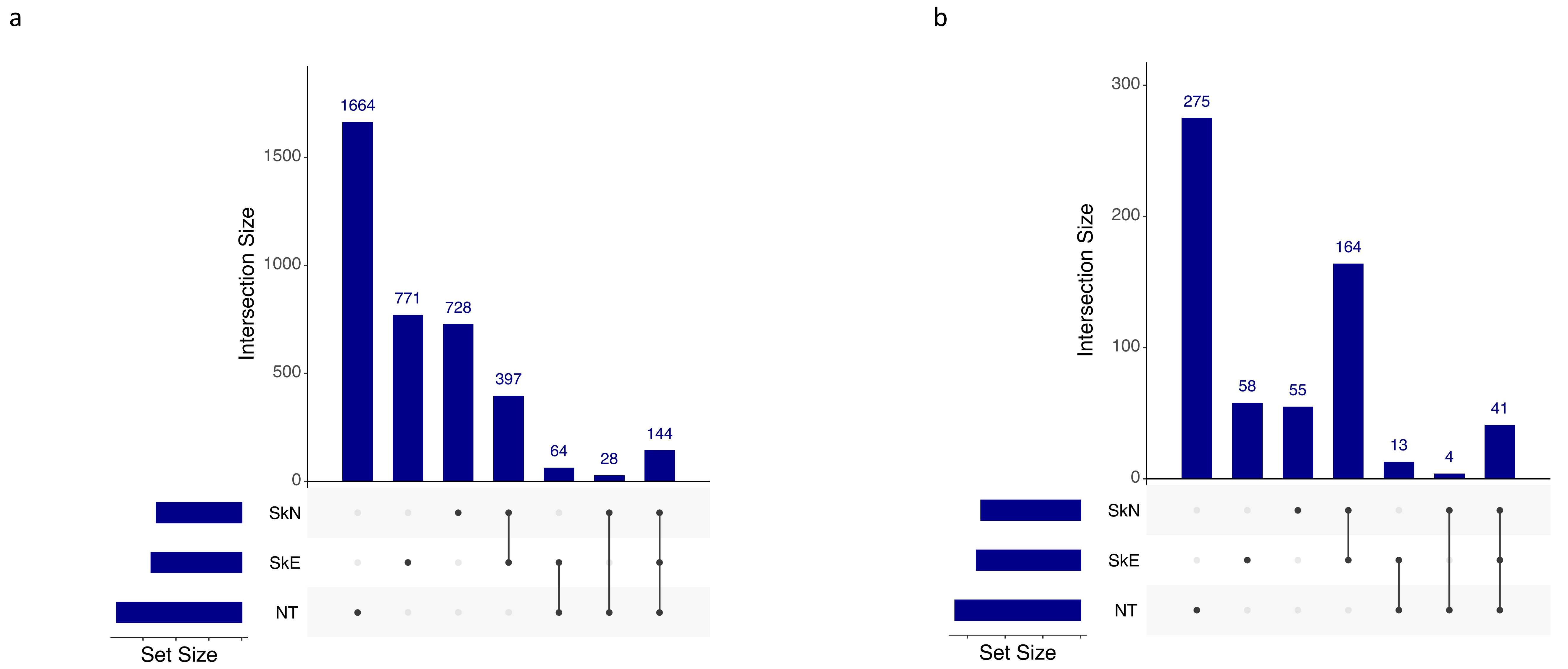


**Supplementary Figure 2.** Relative representation of tissue-specific and shared RsQTL correlations in cis-, defined by (**a**) distance (1e6 nt) and (**b**) co-location within the same gene coordinates. On each graph, the three plots on the left represent exclusive NT, SkE, and SkN, ReQTLs, respectively; the 3-tissue overlapping ReQTLs are shown on the most-right. Higher number of RsQTLs are computed in the nerve as compared to the skin, and higher overlap between the two skin tissues (as compared to between skin and nerve) .
