## Supplementary material for "RsQTL: correlation of expressed SNVs with splicing using RNA-sequencing data": S_Figure_3

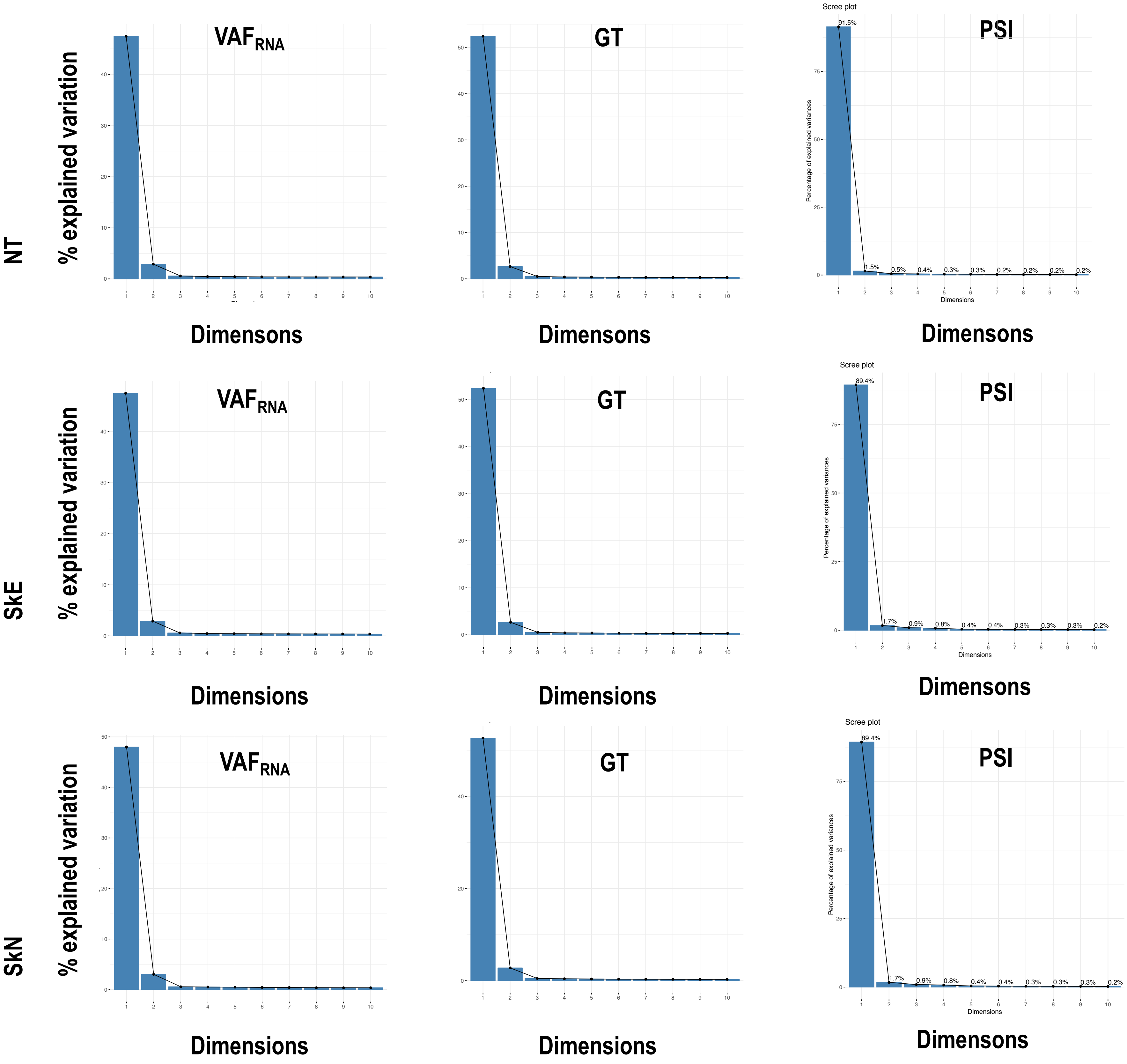


**Supplementary_Figure_3.** Percentage explained variance of the top 10 Principal Components (PC) for VAF$_{RNA}$ Genotypes, and PSI. PCA shows that more than 50% of VAF$_{RNA}$ , GT and PSI variance is explained by the top two PC.
