## Supplementary material for "RsQTL: correlation of expressed SNVs with splicing using RNA-sequencing data": S_Table_2

**Supplementary_Table 2.** Total and shared number of cis-RsQTLs and cis-sQTLs identified in each tissue.

| Tissue | Correlations | | Shared RsQTLs-sQTLs | | |
|---|---|---|---|---|---|
| | RsQTL | sQTL | Total N | % RsQTL | % sQTL |
| **NT** | 1039 | 1620 | 907 | 87.3 | 56.0 |
| **SkE** | 777 | 1231 | 702 | 90.3 | 57.0 |
| **SkN** | 708 | 1166 | 627 | 88.6 | 53.8 |
| **Genes** | | | | | |
| **NT** | 220 | 219 | 169 | 76.8 | 77.2 |
| **SkE** | 161 | 162 | 118 | 73.3 | 72.8 |
| **SkN** | 160 | 185 | 127 | 79.4 | 68.6 |
| **SNVs** | | | | | |
| **NT** | 605 | 922 | 529 | 87.4 | 57.4 |
| **SkE** | 433 | 673 | 385 | 88.9 | 57.2 |
| **SkN** | 388 | 626 | 343 | 88.4 | 54.8 |
